# Timing and Duration of *Gbx2* Expression Delineates Thalamocortical and Dopaminergic Medial Forebrain Bundle Circuitry

**DOI:** 10.1101/579664

**Authors:** Elizabeth Normand, Catherine Browning, Mark Zervas

**Affiliations:** Department of Neuroscience, Cell Biology and Biochemistry, Division of Biology and Medicine, Brown University, Providence, Rhode Island, USA; Department of Molecular Biology, Cell Biology and Biochemistry, Division of Biology and Medicine, Brown University, Providence, Rhode Island, USA

## Abstract

Gene expression is a dynamic process, which is highly coordinated during development to ensure the proper allocation and identity of neuronal cell types within the brain. Equally important during neurodevelopment is how cohorts of neurons establish axonal projections that innervate terminal target sites. We sought to bridge the temporal dynamics of gene expression, within a specific genetic lineage, to the establishment of neuronal circuits derived from cohorts of the lineage-specific progenitors. A central goal was to be able to accomplish genetic inducible circuit mapping non-invasively and with commonly available CreER/*loxP* technology. Specifically, we genetically marked thalamic neuron progenitors that expressed the transcription factor *Gbx2* at an early embryonic stage and tracked the formation of lineage-derived thalamocortical axons during embryogenesis. We then assessed the neural circuitry at an early postnatal stage. We show that the temporal specificity of lineage marking provides a high degree of clarity for following neural circuit development. We also determined that the onset and duration of gene expression can delineate subsets of neural circuits derived from a common lineage. For example, we uncovered a novel contribution of *Gbx2*-expressing progenitors to midbrain dopamine neurons and dopaminergic axons of the medial forebrain bundle. We anticipate that this system can be instructive in elucidating changes in neural circuit development in both normal development and in mutant mice in which neural circuit formation is altered.

## INTRODUCTION

The relationship between a specific gene expressed in progenitors during development and the terminal cell fate of these progenitors is referred to as a genetic lineage. An important aspect of genetic lineage is how the timing of gene expression within the lineage helps shape the distribution and ultimate cell fate in mature tissues including the nervous system [1,2].

An important component of the genetic lineage of neurons and implicit in nervous system development is that axons derived from subsets of neurons anatomically bind discrete functional domains through the establishment of neural circuits. An important developmental problem is forging a link between progenitors that express genes with temporal precision and establishing neural circuits related to the lineage-derived neurons. There have been recent advances in tackling the problem of neural circuit formation using mice as a model system (reviewed in [3] and see [4], but there are relatively few examples of marking neural circuits and establishing easily discernible point to point labeling of axonal connections based on temporal gene expression in mice. Thus, a desirable feature of circuitry mapping would be the use of a genetic-based system that can be used to mark progenitors with spatial and temporal control, conferring high fidelity of marking, and that could be readily used to better assess genetic mutant mice. It was previously shown that an inducible *CreER* recombinase-based system and a conditional GFP reporter allele allowed for the detection of lineage-derived axons [5,6]. More recently, a conditional red fluorescent reporter allele was used to mark/track thalamocortical circuits and the innervation of their target sites in control and mutant mice with fine spatial and temporal control [7].

Here, we use a well-characterized *CreER* line [7,8] in combination with commercially available conditional reporter mice to characterize the distinct stages of axonal development of neurons derived from the *Gbx2* lineage. Specifically, we marked and tracked this lineage with respect to developing thalamic neurons and their emerging circuits during *in vivo* development.

A surprising finding was that the *Gbx2* lineage also contributed to midbrain dopamine neurons and to medial forebrain bundle circuitry. Collectively, these results reiterate that a genetics-approach is advantageous for elucidating temporal dynamics of gene expression and refining lineage-specific neural circuits *in vivo*. We anticipate that the ease and robustness of this approach will advance our understanding of neural circuit formation in normal development and in genetic mutant mice.

## MATERIALS AND METHODS

### Mice and Fate Mapping

*Gbx2^CreER^* mice were bred to *R26^tdTomato^* reporter mice to generate *Gbx2^CreER-ires-eGFP^;R26^tdTomato^* mice that were subsequently bred to Swiss Webster female mice (Taconic) for <u>g</u>enetic <u>i</u>nducible <u>f</u>ate <u>m</u>apping (GIFM) experiments. *Gbx2^CreER-ires-eGFP^* mice [8] were generously provided by James Li (UCHC) while the *Rosa26^tdTomato^* (*Rosa26^lox-STOP-lox-tdTomato^*) [9] line was purchased from Jackson Laboratories.

Adult *Gbx2^CreER-ires-eGFP^;R26^tdTomato^* male mice between four and six weeks of age were set up in breeding pairs with Swiss Webster (Taconic) females and were checked for the presence of a vaginal plug each morning at 0900 hours. The presence of a plug indicated a successful breeding event and was considered 0.5 days *post conceptus* or embryonic day (E)0.5. Pregnant female mice harboring embryos at E9.5 were administered 4mg (200μl) of tamoxifen (T-5648, Sigma) from a stock solution of tamoxifen in corn oil (20mg/ml) at 0900 hours by oral gavage as previously described [5,10,11]. Litters were then dissected at E12.5 or E18.5. An additional cohort of mice that were allowed to go to term were sacrificed at postnatal day (P)7. All mice were housed, handled, and euthanized in accordance with IACUC guidelines at Brown University.

### Tissue Processing

Embryos were dissected in PBS over ice and a small tail biopsy was used for genotyping. P7 mice were perfused with 4% paraformaldehyde (PFA)/saline and brains were removed as previously described [5,10]. Embryos and brains were fixed in PFA overnight at 4°C. Subsequently, tissue was rinsed in PBS, immersed in 15% sucrose, and 30% sucrose until fully submerged. Tissues were embedded in <u>O</u>ptimal <u>C</u>utting <u>T</u>emperature (OCT) media in cryomolds. The OCT/cryomolds were then immersed in a polypropylene beaker containing 2-methyl-butane that was immersed in a vessel containing liquid nitrogen until the temperature reached −150°C as previously described [10]. Sections were obtained with a Leica cryostat and mounted on slides.

### Immunocytochemistry (ICC) and microscopy

Sections (12μm) were rinsed in PBS for 5 minutes (min) and fixed in 4% PFA in PBS for 5 min. Slides were then rinsed 3 times in 0.2% TritonX-100 in PBS (PBT) for 5 min each and blocked in 10% donkey serum in PBT for 2 hours (h) at room temperature in a humid box.

Sections were immunolabeled using an anti-DsRed antibody that detects the protein product generated by the recombined *R26^tdTomato^* reporter allele (anti-dsRed Ab from Clontech, Catalog (Cat) #632496, 1:500 in 10% donkey serum in PBT). We also used an anti-TH primary antibody (Chemicon; Billerica, MA; Cat #AB152, 1:500 in 10% donkey serum in PBT) to detect dopamine neurons and their axons at E12.5. Appropriate, species matched Alexa secondary antibodies (Molecular Probes) were prepared at a concentration of 1:500 in 1% donkey serum in PBT. Sections were incubated in 300μl of secondary antibody solution for 2h at room temperature, washed with PBT five times for 10 min each, and counterstained with .01% Hoechst 33342 (Molecular Probes; Cat # H-3570) in PBS for 5 min in the dark. Slides were washed two times with PBS for 2 min each, dried and coverslipped.

Low magnification images (Figures 1A, 3A, and 4A) were captured with a Leica MZ16F stereo fluorescent dissecting microscope using PictureFrame software. High magnification images images were collected with a Leica DM6000B epifluorescent microscope using Volocity 5.1 imaging software (Improvision) and were obtained using a motorized stage with a 20x objective. True magnifications are indicated by scale bars in figures. All images were pseudo-colored live as part of the acquisition palette and protocol. Subsequently, imaging data sets were exported to Adobe Photoshop and montages of representative data were generated and presented here.

**Figure 1.**
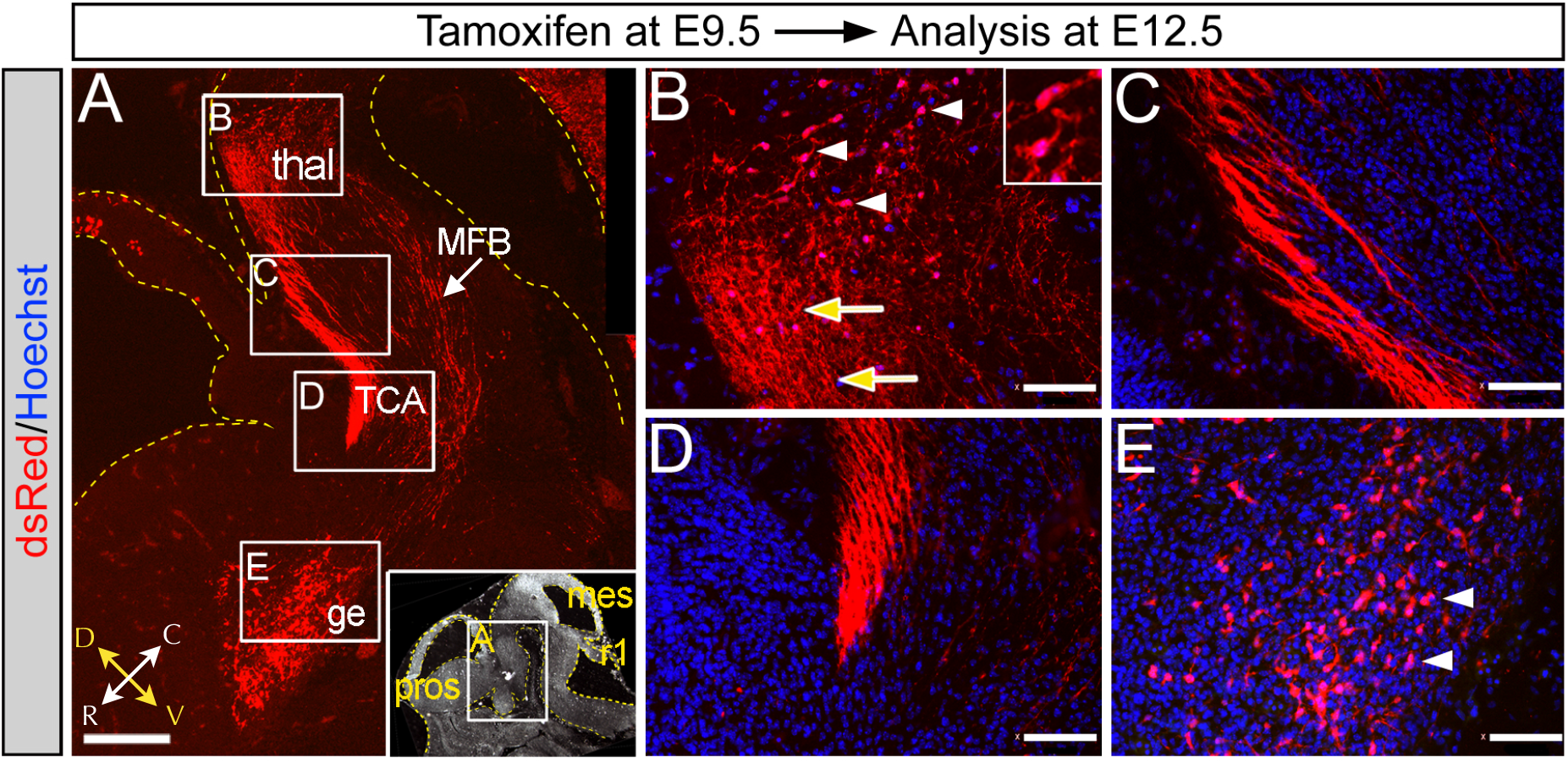
Genetic marking uncovers early neural circuit formation. Tamoxifen was administered to *Gbx2^CreER-ires-eGFP^;R26^tdTomato^* embryos in *utero* at E9.5 and analyzed at E12.5. (**A**) Low magnification sagittal section immunolabeled with an anti-dsRed antibody to detect neurons in the <u>thal</u>amus (thal) and <u>g</u>anglionic <u>e</u>minence (ge) that underwent recombination. Neuronal projections were also labeled with the reporter allele and allowed for the detection of a thick fascicle consistent with <u>t</u>halamo<u>c</u>ortical <u>a</u>xon (TCA) bundles that coursed rostrally and were halted at the ge. A loose axonal bundle was also detected in the vicinity of the <u>m</u>edial <u>f</u>orebrain <u>b</u>undle (MFB). The black and white inset shows regions of analysis and reference structures: prosencephalon (pros), mesencephalon (mes), rhombomere 1 (r1). (**B**) *Gbx2*-derived neurons of the thalamus (dsRed+, arrowheads) and their corresponding thick axonal plexus (arrows). (**C**) TCA projections of *Gbx2*-derived neurons marked at E9.5 exited the rostral thalamus as a thick bundle. (**D**) TCA termination at a site distal to the thalamus. (**E**) *Gbx2*-derived neurons in the ge. In B-E, sections were counterstained with hoechst (blue).

## RESULTS

We administered tamoxifen to E9.5 *Gbx2^CreER-ires-eGFP^* embryos to mark thalamic progenitors *in vivo* [7,8]. We coupled this mouse line with *R26^tdTomato^* conditional reporter mice because they provide a robust readout of Cre-mediated recombination [7,9]. In the absence of tamoxifen, CreER protein is sequestered in the cytoplasm and recombination of conditional alleles does not occur [5,11]. However, delivering tamoxifen to pregnant female mice (see Methods) controls the temporal release of CreER from sequestration, which frees it to translocate to the nucleus where it mediates recombination between same-site oriented *loxP* sites that flank a *stop* cassette in the reporter allele (reviewed in [2]). Thus, tamoxifen administration allows us to control the timing of the recombination event and cell marking within the *Gbx2* lineage.

### *Gbx2* derived neural circuit formation

Subsequent to tamoxifen administration at E9.5, we analyzed embryos at mid-gestation (E12.5) of development, which revealed thalamic neurons that were derived from the *Gbx2* lineage (**Figure 1A,B**). The marked neurons had axons that emerged from the rostral thalamus and formed a thick proximal fascicle, which continued to project rostrally (**Figure 1A-C**). *Gbx2*-derived axonal projections terminated at a distal limit in the proximity of the thalamocortical intermediate target zone (**Figure 1D**) consistent with thalamocortical axon guidance [12]. A second cohort of *Gbx2*-derived neurons were located in the ganglionic eminence (**Figure 1A,E**). In addition to thalamocortical projections, there was a more ventrally located, loose axonal fiber tract in what appeared to be the <u>m</u>edial <u>f</u>orebrain <u>b</u>undle (MFB, **Figure 1A, white arrow**).

### Dynamic gene expression delineates neural circuits

The *Gbx2^CreER-ires-eGFP^;R26t^dTomato^* allelic configuration allows for the determination of progenitors that expressed *Gbx2* at the stage of marking (E9.5, red) and whether they continued to express *Gbx2* at the stage of analysis (E12.5, green). We previously found this approach to be instructive in determining how the timing and duration of *Gbx2* expression shaped spinal cord development [13]. In this study, we assessed dynamic gene expression and early thalamic circuits using lineage analysis and identified neural circuits that were derived from early *Gbx2* expressing progenitors marked at E9.5 (tdTomato+, red) and neurons that continued to express *Gbx2*(GFP) at the time of analysis (GFP+ neurons and their axons, green). Interestingly, the thalamocortical axons (TCA) at E12.5 were derived from neurons that had both early and persistent *Gbx2* expression (tdTomato+/GFP+, yellow) (**Figure 2A,A′**). In contrast, the *Gbx2*-derived axons of the putative MFB did not continue to express *Gbx2*(GFP) (**Figure 2A**). We then used antibody labeling to detect tyrosine hydroxylase (TH) which labels midbrain dopamine neurons and TH+ axons that course along the MFB [14,15]. Surprisingly, *Gbx2*-derived axons (dsRed+) in the MFB were also TH+, which suggests that midbrain dopamine neurons were derived from *Gbx2*-derived progenitors marked at E9.5 (**Figure 2B,B′**). Therefore, we investigated whether midbrain dopamine neurons, are directly derived from *Gbx2*-expressing progenitors.

**Figure 2.**
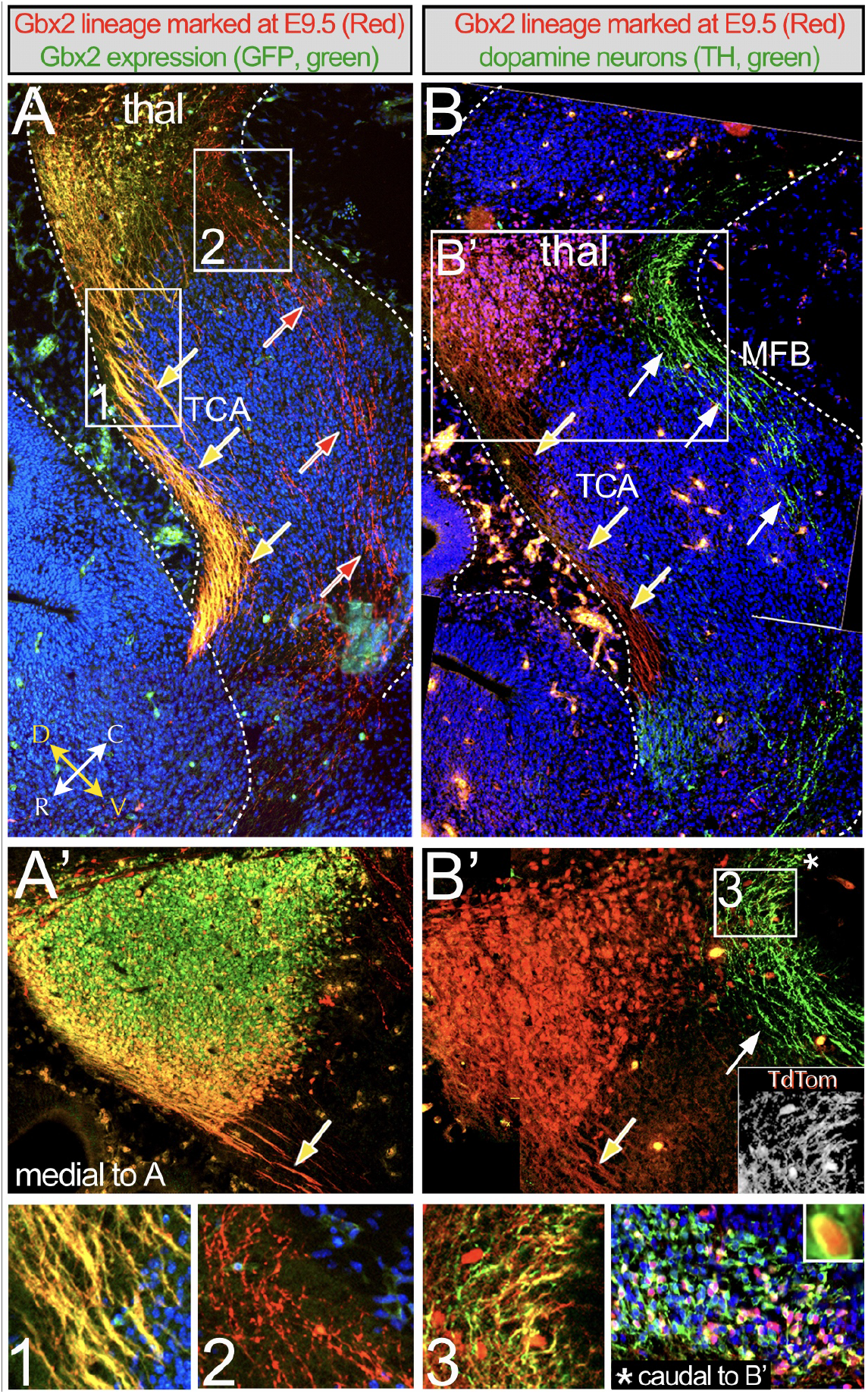
Timing of gene expression defines subsets of neural circuits within a defined genetic lineage. Tamoxifen was administered to *Gbx2^CreER-ires-eGFP^;R26^tdTomato^* embryos at E9.5 and analyzed at E12.5. (**A**) Sagittal section immunolabeled with anti-dsRed and anti-GFP antibodies to detect thalamic neurons that, respectively, expressed *Gbx2* at E9.5 (dsRed+, red) and continued to express *Gbx2* at E12.5 (GFP+, green). Sections were counterstained with hoechst (blue). Neurons in the thalamus (thal) and TCA (yellow arrows) were dsRed+/GFP+ (yellow) indicating that they were derived from neuronal progenitors that expressed *Gbx2* at E9.5 and continued to express *Gbx2* at E12.5. The axons of the medial forebrain bundle (MFB) were dsRed+/GFP− (red arrows) indicating that these projections were related to neurons that expressed *Gbx2* early, but ceased expressing *Gbx2* by E12.5. (**A′**) Thalamus from same sample but at more medial location. (**1,2**) High magnification panels of regions of interest shown in A. (**B**) Sagittal section immunolabeled with anti-dsRed and anti-tyrosine hydroxylase (TH) antibodies to detect neurons that expressed *Gbx2* at E9.5 (dsRed+, red) and to identify dopamine neurons (TH+, green), respectively. Neurons in the thal and TCA were dsRed+ consistent with adjacent section in A. The MFB contained TH+ dopaminergic axons (white arrows). (**B′**) Thalamic and dopaminergic axons coursing ventral to the thal. The TH+ projections were also dsRed+ but were much fainter than the neurons in the thalamus. Black and white inset shows red channel alone from region 3. (**3**) High magnification panel of region 3 with the red signal increased to observe the projections. The asterisk (*) in B′ is a reference for the domain caudal and medial to panels shown in B,B′ at the ventral mesencephalic flexure. The domain is depicted in the panel [*caudal to B′] and shows midbrain dopamine neurons derived from the *Gbx2* lineage marked at E9.5 (dsRed+/TH+, yellow) that project rostrally via the MFB.

It was notable that co-immunolabelling with TH and dsRed antibodies validated that some midbrain dopamine neurons (TH+) were derived from the *Gbx2* lineage (dsRed+). During embryogenesis *Gbx2-* expressing progenitors are positioned caudal to the ventral mesencephalon and located in the ventral domain of rhombomere 1 [16]. It was previously shown that a lineage restriction boundary is established between the ventral domains of the mesencephalon and r1 shortly after E9.5 [17], which prevents midbrain progenitors from contributing to r1-derived structures. However, these findings suggest that the boundary is not bi-directional and that some ventral r1 progenitors expressing *Gbx2* traverse this (mesencephalic) lineage boundary and contribute to midbrain dopamine neurons (**Figure 2B′** and the inset \***caudal to B′**). Thus, both TCA and MFB circuits are derived from the *Gbx2* lineage at E9.5, but the TCA bundle persists in expressing *Gbx2* at least until E12.5 while the MFB only transiently expresses *Gbx2*, which is ceased by E12.5.

### Maturation of thalamocortical circuits

The *Gbx2* lineage-derived neurons marked at E9.5 continued to populate the thalamus at E18.5 (**Figure 3A,B**). These thalamic neurons had axons that exited the ventral aspect of thalamus (**Figure 3A,C**) and traversed ventrally and rostrally as fascicles, which passed through the ventral half of the striatum (**Figure 3A,D**). The bundles de-fasciculated as they entered the cerebral cortex and formed fine axonal branches, which traversed the deep cortical layers (**Figure 3A,E**). In addition, tdTomato (dsRed+) projections were used to follow entire axonal bundles oriented in parallel to each other *en route* to the deep cortical layers in more lateral sections, which revealed robust innervation of layer 6 (**Figure 3F**). Projections proximal to layer 6 were dense (in deep layers), but without obvious orientation. The marked projections became much sparser in more superficial cortical layers (**Figure 3F**).

**Figure 3.**
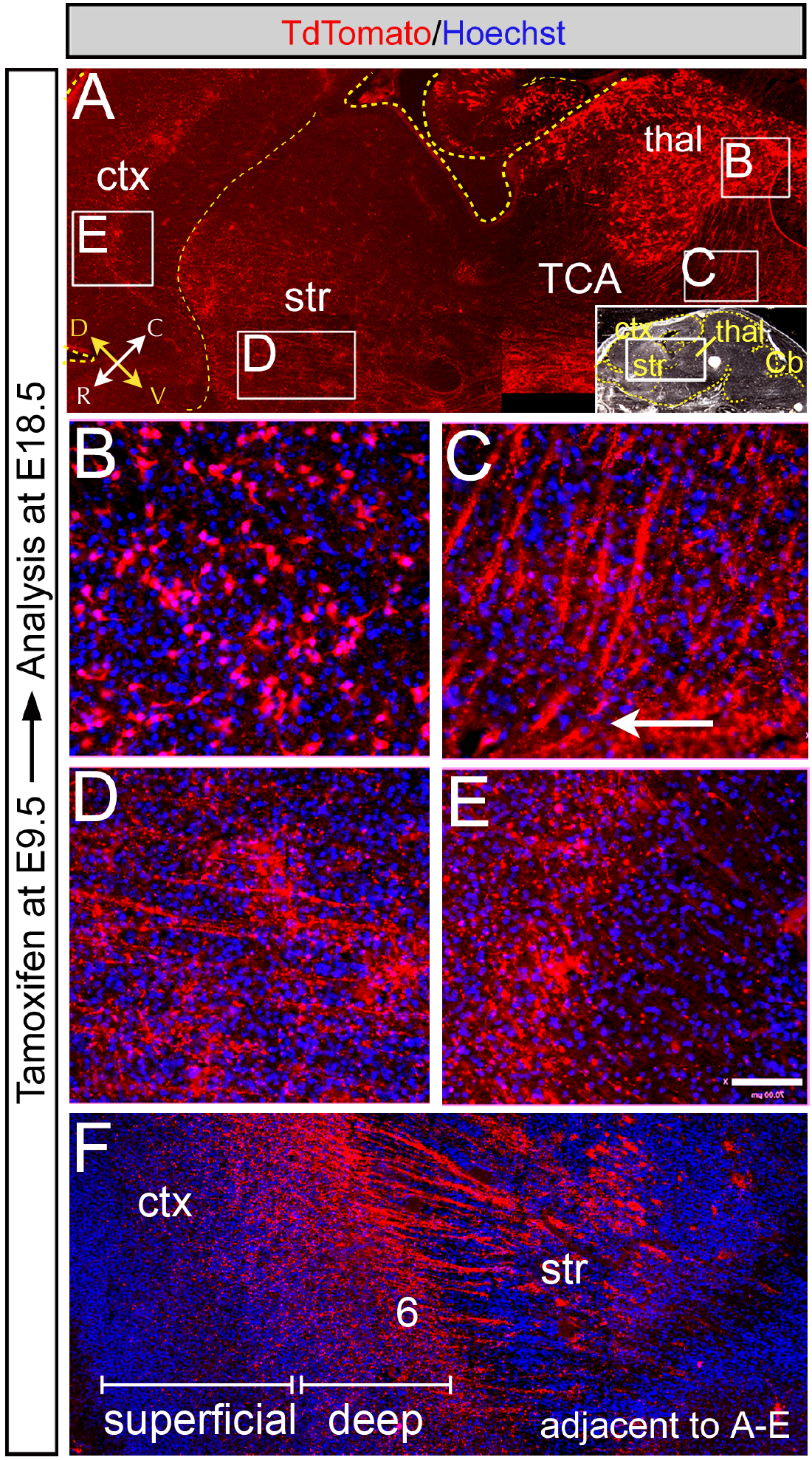
Maturation of lineage derived neural circuits. Tamoxifen was administered to *Gbx2^CreER-ires-eGFP^;R26^tdTomato^* embryos at E9.5 and analyzed at E18.5. (**A**) Sagittal section immunolabeled with an anti-dsRed antibody to detect neurons in the thalamus (thal) that underwent recombination. Sections were counterstained with hoechst (blue). The TCA bundle traversed the striatum (str) and entered the deep layers of the cerebral cortex (ctx). (**B**) Thalamic neurons that expressed *Gbx2* at E9.5 were becoming morphologically distinct. (**C**) The TCA fascicles exited the thal as thick well-defined processes that turned rostrally (arrow). (**D**) The *Gbx2* derived projections entered the striatum and formed a fine network of axon collaterals, which was in contrast to TCA marked at the same stage, but analyzed at E12.5 (See Figure 1). (**E**) TCA that exited the striatum ramified the the deep cortical layer. (**F**) In a section adjacent to panels A-E, *Gbx2*-derived TCA passed through the striatum and innervated deep cortical layers with fine axonal branches that were progressively sparser as they ramified layers more superficial to cortical layer 6.

### Early postnatal thalamocortical circuits

Finally, thalamic neurons derived from *Gbx2*-expressing progenitors marked at E9.5 were distributed broadly throughout the thalamus by the first postnatal week (P7) (**Figure 4A,B**). These neurons had characteristic thalamic morphologies (**Figure 4B, inset**). In addition, thalamic neurons had broad sweeping fascicles that apparently emerged from the rostral-lateral extent of the thalamus and entered the internal capsule (**Figure 4A,C**).

**Figure 4.**
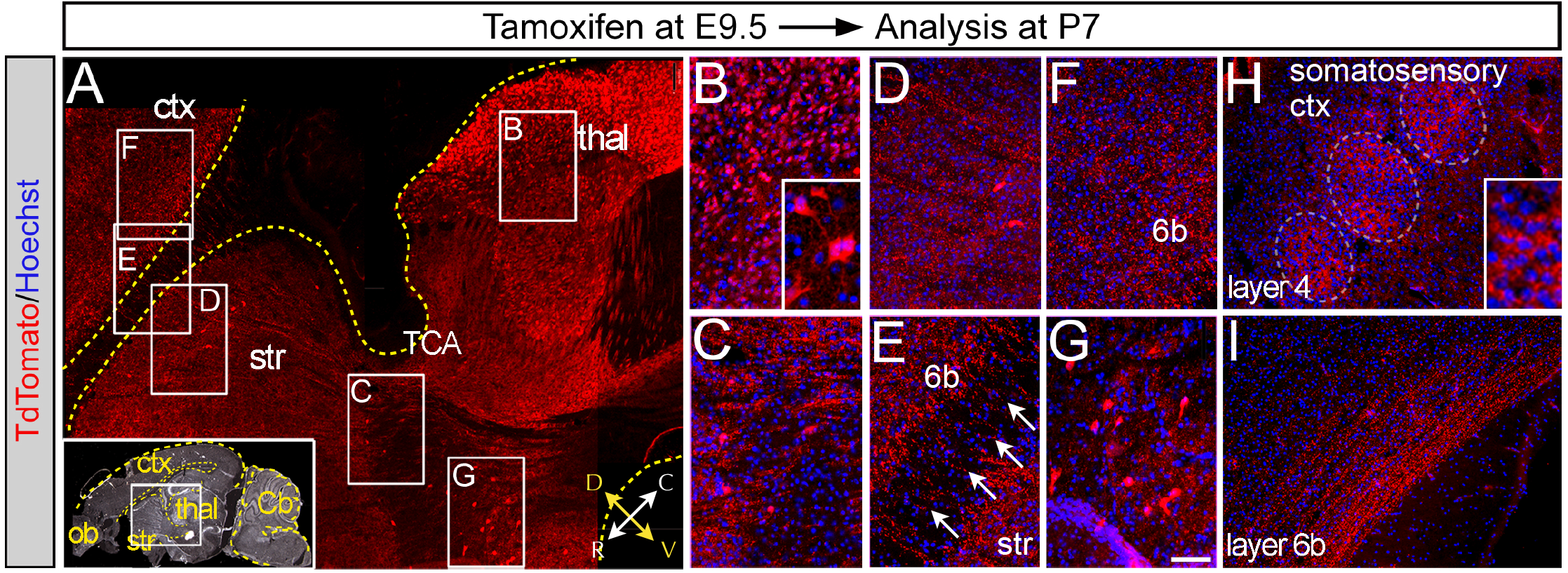
Target innervation of genetic lineage derived neural circuits. Tamoxifen was administered to *Gbx2^CreER-ires-eGFP^;R26t^dTomato^* embryos at E9.5 and analyzed at P7. (**A**) Sagittal section immunolabeled with an anti-dsRed antibody detected neurons and the thalamocortical axon bundle (TCA) derived from the *Gbx2* lineage marked at E9.5. Sections were counterstained with hoechst (blue). *Gbx2*-derived TCA traversed the striatum (str) as thick bundles where they branched and formed fine axonal terminations. The TCA exited striatum and innervated the cerebral cortex (ctx). (**B**) Neuronal progenitors that expressed *Gbx2* at E9.5 contributed to mature thalamic neurons that were distributed in the thal. (**C**) Numerous thick TCA bundles exited the thal. (**D**) TCA traversed the entire distance of the str as thick bundles with fine axonal branches that established a *Gbx2*-derived plexus in the caudate/putamen of the str. (**E,F**). Upon exiting the str, the TCA projections were thinner in comparison to when they entered and traversed the str. The TCA then formed a a dense plexus in deep cortical layer 6b. (**G**) *Gbx2*-derived neurons remaining in close proximity, but ventral to the thal. (**H,I**). In an adjacent sagittal sections to A-G, *Gbx2*-derived axons innervated somatosensory cortex where the axons coalesced in a fingerprint pattern, consistent with the early formation of barrel cortex (H). In addition, *Gbx2*-derived axons traversed the rostral-caudal axis as long axonal bundles that were oriented parallel to the ventral surface (I).

The fascicles remained well organized and appeared as parallel-oriented fibers with numerous small terminal ramifications throughout and at the anterior-dorsal extent of the caudate/putamen (**Figure 4A,D**). Significant cohorts of thalamocortical projections passed through the internal capsule and caudate/putamen as smaller bundles when compared to their entrance into the internal capsule. These *Gbx2*-derived projections retained a parallel orientation with each other as they entered into layer 6b of the cerebral cortex (**Figure 4A,E,F**).

*Gbx2*-derived neurons were also located ventral and rostral to the thalamus (**Figure 4A,G**). In adjacent parallel sections (**Figure 4H,I**) the *Gbx2*-derived axons innervated somatosensory cortex resulting in the coalesced innervation of the immature somatosensory barrels of cortical layer 4 (**Figure 4H, inset**). In addition, deep-layer projections were organized as a dense fiber tract with long, longitudinal processes orientated along the rostral-caudal axis within layer 6b (**Figure 4I**). In summary, aside from somatosensory innervation, once thalamocortical axons innervated the cerebral cortex, these fibers de-fasciculated and formed numerous fine ramifications in the cortex. However, in somatosensory cortex, *Gbx2*-derived projections formed dense axonal clusters and established the rudimentary somatosensory barrels.

## DISCUSSION

The thalamus is comprised of both locally projecting interneurons and long-distance projection neurons [18]. Physical tracing methods have been instructive in elucidating the formation of thalamocortical axons and is a well-described procedure that shows that thalamic axons emanate from the thalamus and halts at intermediate target sites. Subsequently, thalamic axons turn toward and innervate the cerebral cortex [12,19,20,21]. It has been shown that the *Gbx2* lineage contributes to thalamic neurons and that *Gbx2* is required for the proper formation of thalamic axonal projections [7,22,23]. In addition, *Gbx2* is required cell non-autonomously for thalamic development [8].

However, the methods used in the aforementioned thalamocortical circuitry studies have been invasive and, therefore, may have labelled fibers of passage. Additionally, a relationship between the timing of gene expression, genetic lineage, and neural circuit formation has not been established for the *Gbx2* lineage. Thus, we sought to use GIFM as an approach to demonstrate whether the early expression of *Gbx2* in thalamic neuron progenitors was related to the formation of long distance axonal circuits that eventually innervate cortical targets. We marked *Gbx2* expressing progenitors at E9.5 and analyzed early patterns of axonal growth after three days. We observed clearly labeled *Gbx2* lineage-derived thalamocortical axons that exited the thalamus in thick fascicles and were abrogated at the ganglionic eminence consistent with previous physical axonal tracing methods [18]. By E18.5 *Gbx2* derived axons entered the striatum as thick fascicles and formed a rich axonal plexus. Upon exiting the caudate/putamen, well-defined axonal tracts entered the deep cortical layers with fine axons that began to ramify into more superficial layers. At P7, the axons that entered the cortex formed an axonal tract that traversed the rostral-caudal axis of the cortex. Notably, at E18.5 only fine axons innervated more superficial cortical layers, while at P7 the projections establish a broad zone of termination in layer 4 of cortex. In the somatosensory cortex, the projections coalesced as the rudimentary layer 4 barrel structures.

During the course of our analysis, we also observed a second, surprising *Gbx2* lineage-derived axonal tract. This second bundle was located ventrally to the thalamus as part of the putative medial forebrain bundle. Thus, we show for the first time that a portion of medial forebrain bundle axons are derived from *Gbx2*-expressing progenitors that contribute to dopamine neurons. It is well known that *Gbx2* expression defines a domain that is posterior to the germinal zone of ventral midbrain progenitors [24,25,26,27]. However, a recent study showed that *Gbx2* transcript is expressed in the ventral mesencephalic midline at E10.5 and that *Gbx2* lineage contribution occurs in the ventral-medial midbrain primordia [28]. This ventral-medial primordia is the *in vivo* source of midbrain dopamine neurons [29,30], which are connected to the cerebral cortex via the medial forebrain bundle. However, the aforementioned study [28] did not demonstrate a direct link between *Gbx2*-expressing progenitors and terminally fated midbrain dopamine neurons and dopaminergic circuitry. It is quite likely that a portion of the dopaminergic projections that were derived from the *Gbx2* lineage are part of the nigrostriatal bundle in addition to the medial forebrain bundle.

We show here that thalamocortical axons were derived from neuronal progenitors that had early (E9.5) and persistent (E12.5) expression of *Gbx2* (tdTomato+/GFP+, yellow). In contrast, dopaminergic axons of the medial forebrain bundle (and their related dopamine neurons) were derived from progenitors that expressed *Gbx2* only early and transiently (at E9.5, but not at E12.5).

In summary, this genetic inducible circuit mapping approach makes it possible to link the differential timing of gene expression and specific neural circuits within a unique genetic lineage. Finally, we show that a conditional genetic-based approach is highly effective at marking and following neural circuit formation and maturation *in vivo*. This genetic inducible circuit mapping approach has distinct advantages over traditional methods including that it is non-invasive and is highly reproducible. In addition, there is no confounding labeling of fibers of passage. In conclusion, this combination of GIFM and circuit tracing forges a link between temporal gene expression during embryogenesis and postnatal neural circuits.

## ACKNOWLEDGEMENTS

This work was supported by a Department of Defense CDMRP grant (TS110067, MZ) and the Brown University Department of Neuroscience training grant (NS062443-02).

